# The First Reconstruction of Intercellular Interaction Network in *Mus musculus* Immune System

**DOI:** 10.1101/820316

**Authors:** Somayeh Azadian, Javad Zahiri, Seyed Shahriar Arab, Reza Hassan sajedi

## Abstract

Intercellular interactions play an important role in regulating communications of cells with each other. So far, many studies have been done with both experimental and computational approaches in this field. Therefore, in order to investigate and analyze the intercellular interactions, use of network reconstruction has attracted the attention of many researchers recently. The intercellular interaction network was reconstructed using receptor and ligand interaction dataset and gene expression data of the first phase of the immunological genome project. In the reconstructed network, there are 9271 communications between 162 cells which were created through 460 receptor-ligand interactions. The results indicate that cells of hematopoietic lineages use fewer communication pathways for interacting with each other and the most network communications belong to non-hematopoietic stromal cells and macrophages. The results indicated the importance of the communication of stromal cells with immune cells and also high specificity of genes expression in these cells. The stromal cells have the most autocrine communication, and interactions between the wnt5a with the Ror1/2 and Fzd5a among the stromal lineage cells are abundant.

## Introduction

One of the important features of unicellular organisms, and multicellular organisms in particular, is their ability to respond to external stimuli through cellular signaling process. Cellular signaling occurs during two main steps of signal transmission and transduction. In the signal transmission step, first messenger sent from the source of the signal-cells or environment of the target cell-are received by the target cell receptors through transmission in the extracellular matrix. In the signal transduction stage, the receptor-ligand complex, triggers intracellular signaling pathways by secondary messengers and ultimately, by affecting the expression level of related genes, causing cellular responses in the target cell [1]. Cellular signaling involves in adaptation of cell to different conditions, maintaining cellular hemostasis, growth, differentiation, migration, survival, apoptosis, cellular motility, etc. [2]–[7]. In intercellular signaling a cell as the sender of the signal, the signal carrier molecule (first messenger or ligand), an environment through which the messenger molecule is transmitted, and a receptor from signal recipient cell are essential [8][9].

Intercellular communication plays an important role in the behavior of cells and regulates their communications with each other and with the environment, particularly in multicellular beings. In multicellular organisms, cellular behavior is much more complicated than that of a unicellular organism and their ability to communicate with adjacent cells is the basis of their coordinated activity, and formation and maintenance of their integrity [10]. Thus, the necessity of studying different aspects of this process is essential. So far extensive studies have been done with both experimental and computational approaches in this field. Due to the difficulty of studying this processes in a large scale manner in the presence of different types of cells, the use of computational systems biology approaches has recently attracted the attention of many scholars in order to investigate and analyze the intercellular signaling process [11]. Currently, most of the reconstructed networks have been linked to the intracellular signaling networks, and intercellular interactions that trigger the intracellular signaling have been underestimated. In most of the reconstructed networks of intercellular interactions, a limited number of cells have been examined. Only in two studies, an intercellular interconnection network has been investigated in a variety of cells [12], [13]. In this study, the intercellular interaction network between the immune cells and stromal cells of *Mus musculus* is being reconstructed and analyzed.

## Results

After integrating two data sets, based on the threshold defined, from199 cell samples, 162 cells had receptors and ligands that were linked together through 9271 communication paths. From these cells, 102 cells were both sender and recipient of the signal, while 14 cells were sender and 46 cells were recipient of signals. As shown in *Figure1a*, more than half of the interactions between the receptors and the ligands were removed, and this volume of communications on the network was only created by 460 interactions between 202 receptors and 226 ligands.

### • Non-hematopoietic stromal lineage has the highest number of expressed receptor and ligand genes among other lineages

In network, the non-hematopoietic stromal lineage expresses the largest number of receptor and ligand genes. Among the immune lineages, the macrophage lineage has the highest expression of receptor and ligand genes compared to other immune lineages and NK cells have the lowest amount of gene expression with expression of 3 receptor genes and 5 ligand genes *(Fig. 1b)*. The results of this reconstruction indicate that cells belonging to the immune systems use less pathways to communicate with each other than non-hematopoietic cells that these results confirm the results of previous study in this [12].

**Figure 1:**
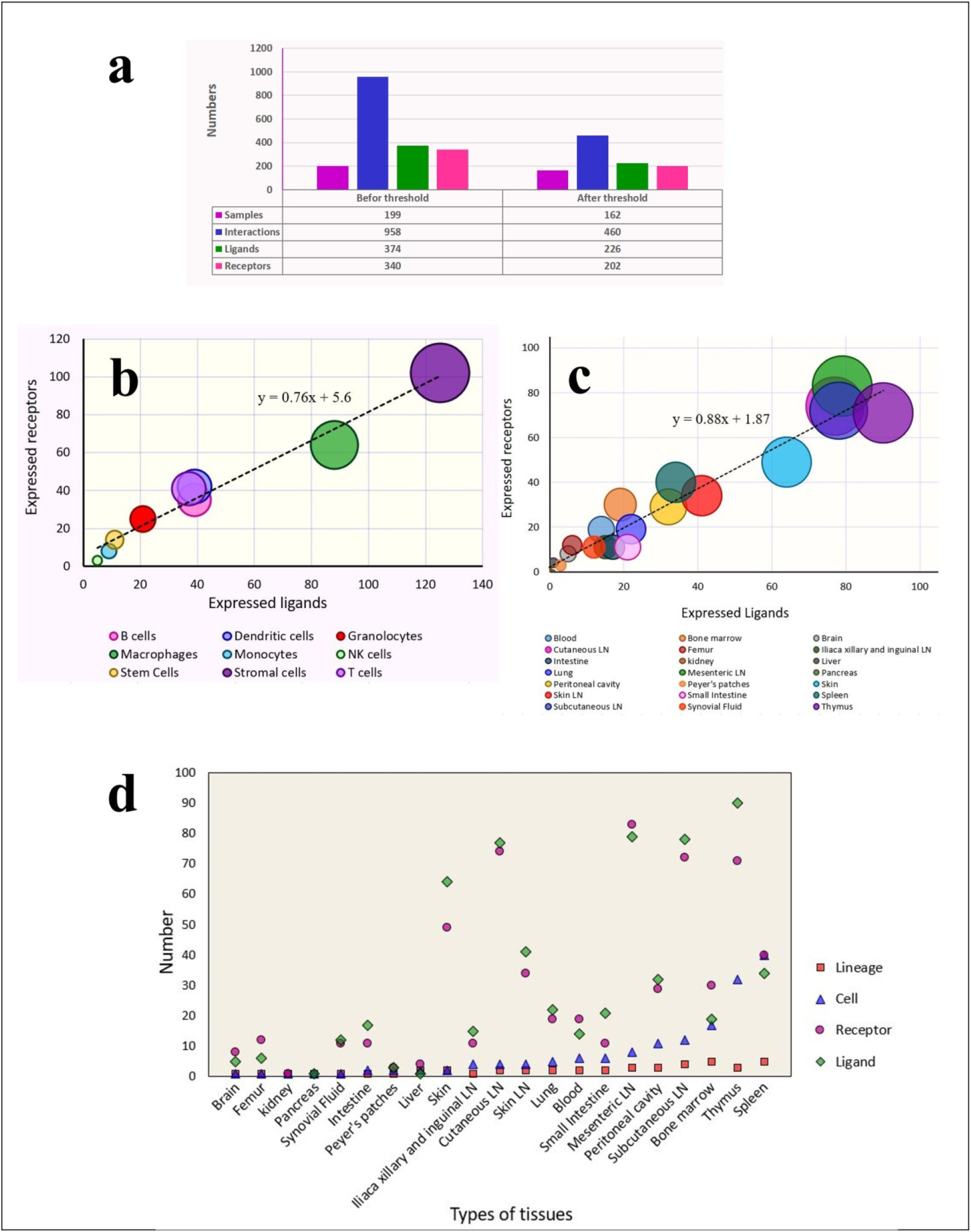
The results of network reconstruction. **a)** Number of cells, number of interactions and number of receptors and ligands involved in the interactions before and after the threshold **b)** Level of expression of receptor and ligand genes in different lineages of the network **c)** Level of expression of these genes tissues **d)** Number of cells, lineages, and the expressed receptors and ligands in each tissue. In figures of 1b and 1c size of each circle represents the sum of the expressed receptors and ligands in that lineage and source.

### • Thymus tissue has the highest number of expressed receptor and ligand genes among other tissues

Examination of the tissue distribution of the cells and expressed ligands and receptors in each cell in the network is shown mesenteric and thymic cells with expression of 162 genes (79 ligand genes and 83 receptors genes) and 173 genes (90 ligand genes and 71 receptors genes) respectively, had the highest expression of ligands and receptor compared to other tissues (*Fig. 1c*). In this network, cells of kidney, and pancreas tissue exhibited the least expression receptor and ligand genes respectively, compared to other tissues. The number of cells, lineages, and also the number of receptors and ligands expressed in each tissue are shown in *Figure1d*. The results are shown tissues with more number of cells do not necessarily have more receptors and ligands for example in the spleen with 40 types of cells, the number of receptors and ligands is significantly less than the thymus gland with 32 types of cells.

This can be due to the different type of lineages that exist in each of these two tissues. In spleen tissue there is five lineages of B lymphocytes, dendritic cells, macrophages, natural killer cells and T lymphocytes, whereas the lineages of thymus tissue are including the T lymphocyte, dendritic cell, and stromal cell lineages. Therefore, in the tissues that there is stromal cells lineage (skin, thymus gland, skin lymph nodes, subcutaneous lymph nodes and mesentery), the expression of the receptors and ligands is much higher than tissues without them.

### • There is a correlation between the frequency of receptors and ligands, and their degree of specificity

The investigation of the cell type-specific expression of genes showed that 52 ligands and 45 receptors express only in one cell type and therefore have specificity degree of 1. The Ccl2 ligand and the Ccr1 receptor, with expression in 15 and 14 cells, respectively, had the lowest degree of specificity compared to other receptors and ligands. The assessment of ligands and receptors expressed in each specificity degrees (Fig. 2a and 2b) indicates the high specificity of the expression of the genes. Therefore, a decreasing trend could be seen in the number of expressed ligands and receptors simultaneously with the decrease in the level of specificity. The survey of ligands and receptors that have the most specificity degree indicates that these genes are expressed in 21 and 29 different cell types respectively (Fig. 2c and 2d) and thus some cells express more than a ligand and receptor with a specificity degree of one. In between these cells, receptors and ligands with high specificity degree are expressed in stromal cells and macrophages.

**Figure 2:**
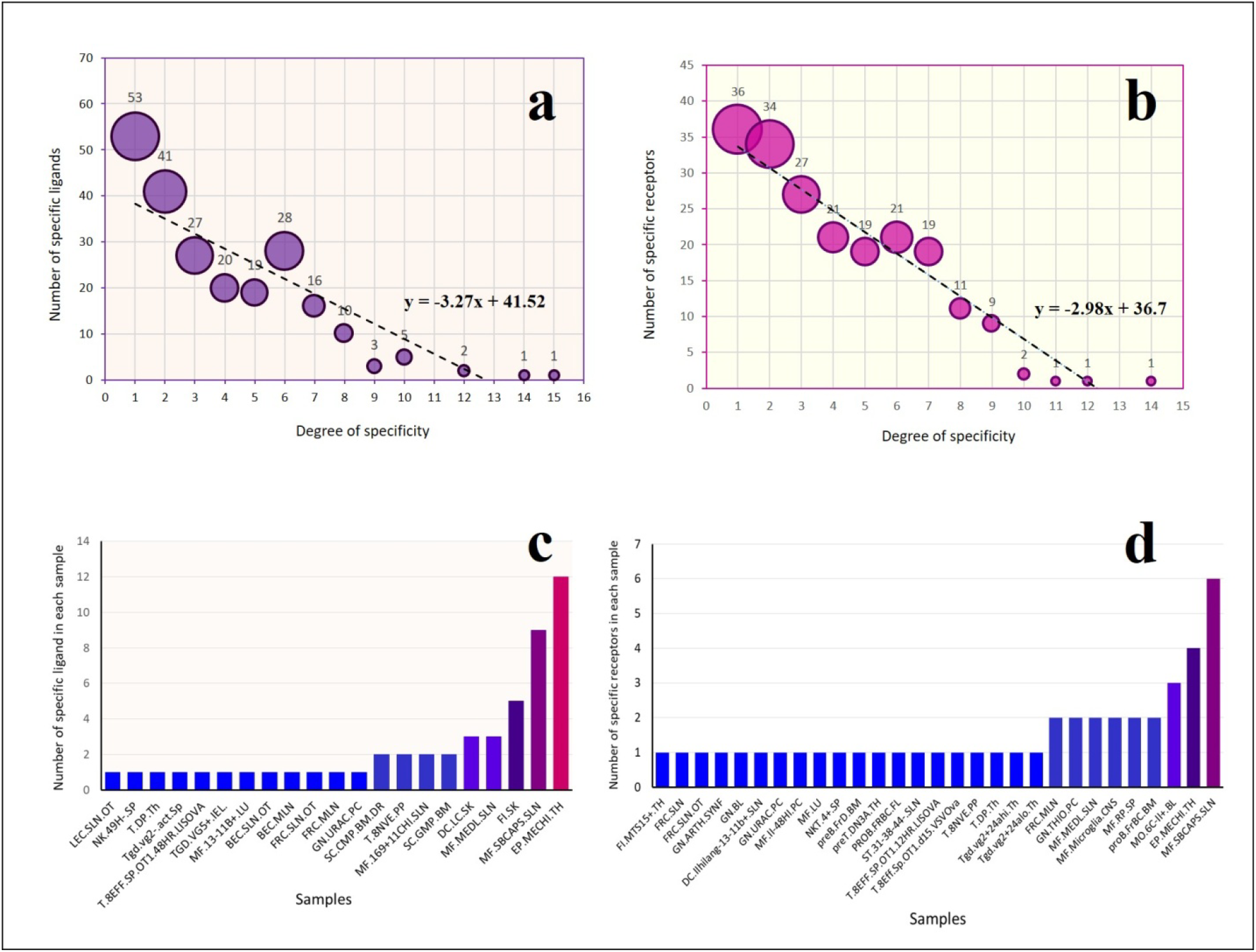
Specificity of cell expression of receptors and ligands **(a** and **b)**.The number of receptors and ligands with the highest specificity in each cell type **(c** and **d)**:

The results obtained from reviewing the specificity of the interactions and the 9271 communication in the network were shown in *Figure 3*. The distribution of the interaction specificity between receptor and ligand among 162 cells has been investigated, which indicates the high specificity of these interactions. Interactions with the specificity degree of 15 for ligand and 14 for receptor are interactions with lowest specificity degree, whereas the lowest degree of interaction specificity that could exist in this network with 162 cells is the interaction with specificity degree of 162 for both receptor and ligand.

**Figure 3:**
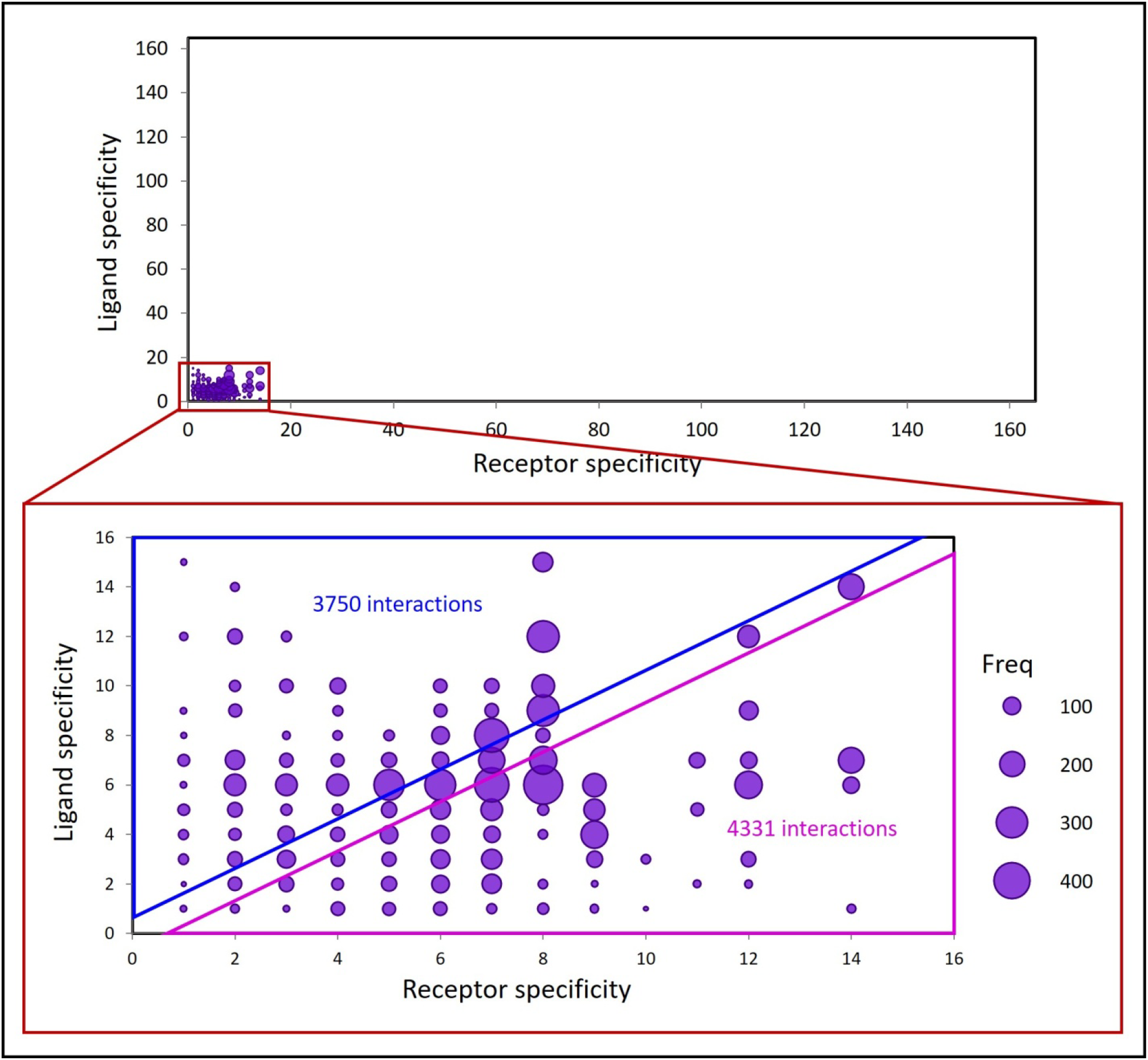
Specificity of the receptor-ligand interactions: The frequency of each specificity degree is determined by size.

The frequency of interactions occurring at each specificity degree are shown in Figure 3, specificity of the interactions is between 1 to 14 for the receptors and 1 to 15 for the ligands therefore the interactions have specificity of 1-1 to 14-15.The interactions with specificity of 6 for the ligands, and 8 for the receptors (6→8) have the highest frequency in network with 432 interactions, and the lowest frequency is related to the specificity of 1→10 and 2→1.Among the9271 interactions, the frequency of interactions in which the receptor specificity is lower than the ligand is 4331, for interactions in which the specificity of the ligand is less than the receptor is 3750, and in 1190 interactions the specificity of the receptor and the ligand are equal.

### • Stromal cells have the highest number of communications in the network

The non-hematopoietic stromal cell lineage has the most communications in network and also among the hematopoietic lineages, B lymphocyte have the most communications. The T lymphocytes in this network have the least communication with other cells. Two stromal cells FRC.SLN and FRC.MLN respectively with output node degree of 651 and input degree of 381 have the most communication as the sender and as the recipient of the signal in the network. Stromal cells associated with the thymus, lymph nodes and subcutaneous lymph nodes have the highest degree of closeness centrality.

### • Stromal cells have the most autocrine communication

In order to determine what cells are involved in autocrine communications, the network was examined based on the presence or absence of the loop (*Fig.4a*). The stromal cells in this network have the most autocrine communications in comparison with the other cells. In *Figure4b*, the cells were presented based on their lineage. Non-hematopoietic stromal cells have the most communications with themselves and with other cells. Although the stem cells are more than monocyte lineage cells but have the smallest amount of communications in the network and have a very small share of the network communications. The intercellular communication network based on their tissues was shown in *Figure 4c*.

**Figure4:**
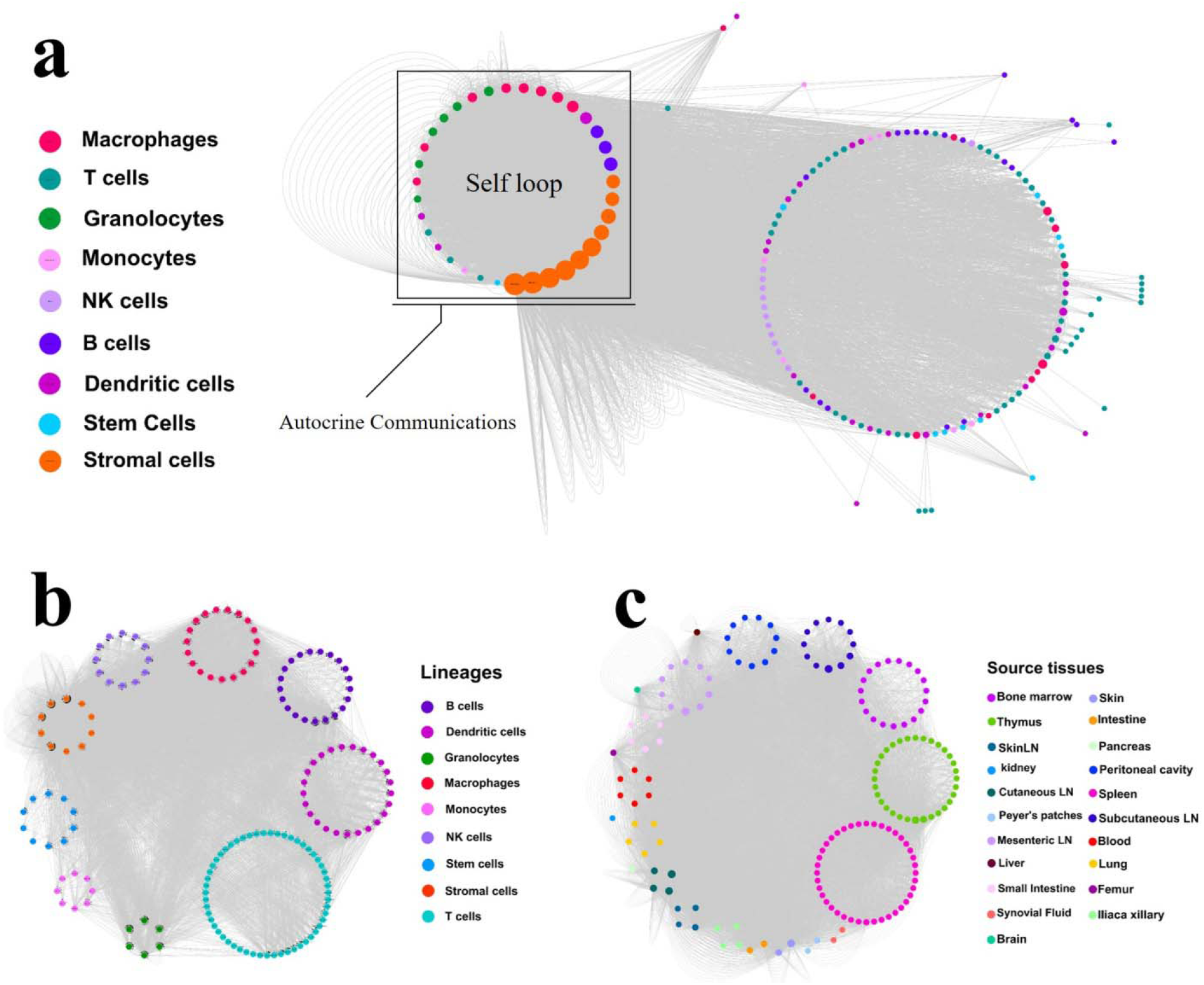
Intercellular communications network. **a)** Intercellular interaction network, determining the existence of autocrine interactions. **b)** Intercellular interaction network represented by different lineages **c)** and different tissues.

### • Macrophages have the most intra-lineage communications compared to other immune lineages

In the inter-lineage interaction network visualized by the Cytoscape (Fig. 5a), there are 9 cell lineages. The linking edges between the lineages are weighted, which indicates the difference in the number of signals that are exchanged between different network lineages. In this network, lineages are directly related to each other therefore the parameter of betweenness centrality of these nodes is low. In this network, the parameter of the clustering coefficient calculated for all nodes is high because neighboring lineages of each lineage interact with each other. The stromal cell lineage has the most intra-lineage communications in comparison to other lineages. The NK and stem cells lineage have no intra-lineage communications. In this network, the stromal cell lineage sends and receives the most signals, and the NK and monocyte lineage sends and receives the least signals compared to other lineages respectively (Fig. 5b).

**Figure5:**
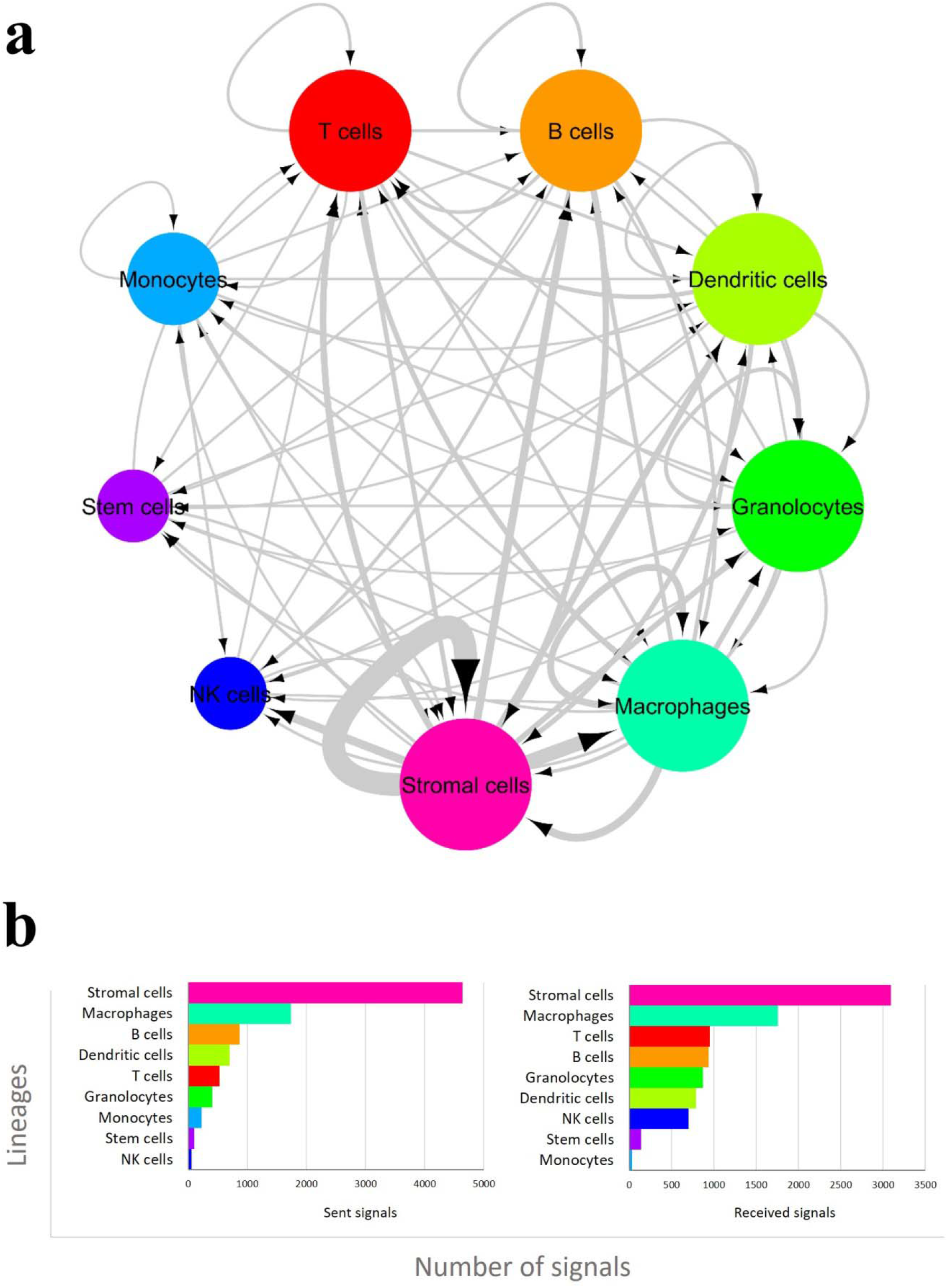
Inter-lineage communication network. **a)** Lineages were sorted based of degree distribution. **b)** Number of sent and received signals in each lineage.

In visualization of the intercellular communication network with a three component pattern in which besides cells, receptors and ligands are also displayed as network nodes, more information can be displayed visually from the network [1].The number of cells that only send the message is less than the other cells, and the cells that act on the network as the sender and recipient of the signal are more abundant than the rest of the cells. Integrin receptors and also chemokine receptors have the highest degree of communication with the ligand molecules *(Fig. 6)*.

**Figure 6:**
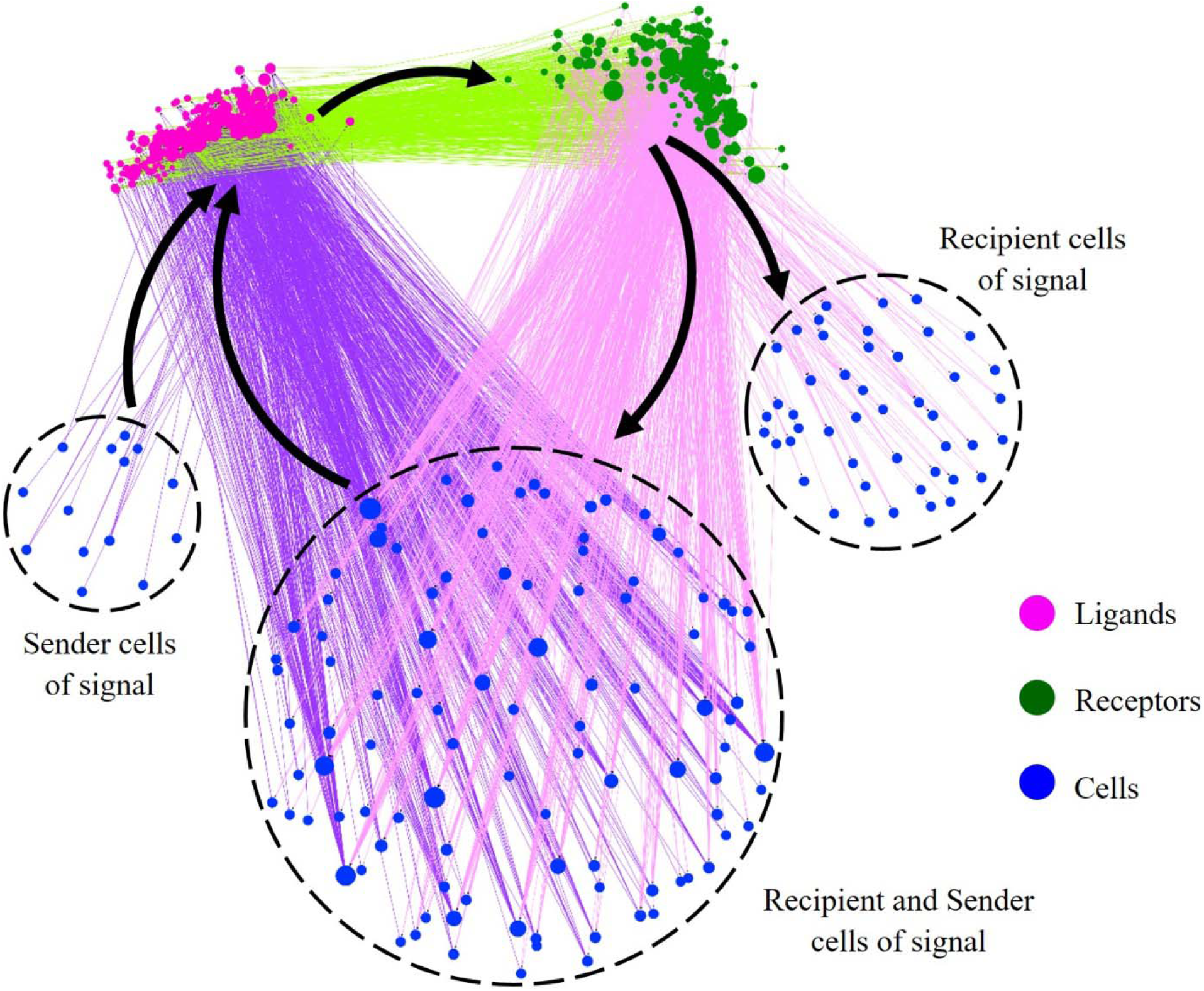
Visualization of the intercellular interactions tripartite network.

## Discussion

In this study, a large number of receptor and ligand genes were expressed in stromal cells and macrophages. This suggests that the type of cell involved in cellular communication has a great influence on the level of the expression of these genes. Stromal cells are non-hematopoietic stem cells that are present in most organs of the body, such as skin. Kidney, pancreas, bone marrow, lymph nodes, and the like. Tissue -resident stromal cells play an important role in the differentiation and maturation of the tissues cells and their progenitors [14].These cells have a supporting role in the function of cells and also participate in hematopoiesis and its regulation[15]. One of the reasons for the high expression of receptor and ligand genes in these cells can be due to different roles that play in each body organ, which requires expression of different types of receptor and ligand molecules to interact with cells located in each organ. In the lymph nodes (LNs), the interactions between the stromal cells, T cells, B cells and dendritic cells lead to the initiation and maintenance of the adaptive immune response process, and on the other hand, these cells also organize the function of the lymph nodes [16], [17].

Researches have shown that markers of skin-resident stromal cells are very similar to markers of mesenchymal stem cells and can be differentiated to adipogenic and osteogenic cells [18]. T cell development depends on a variety of thymic stromal cells which are located in a specialized microenvironment[19]In this network, receptor and ligand genes of stromal cells are expressed in thymus, skin and lymph nodes. Most of the ligands synthesized by stromal cells are cytokines, growth factors, and ECM proteins, and the receptors expressed by these cells also include RTK and RSTK receptors which categorized as enzyme-linked receptors. Gene enrichment analysis of ligands and receptors expressed of stromal cells that were carried out by the DAVID tool [20] shows these genes are present in biological processes such as the development of multicellular organisms, tissues and organs development, positive regulation of cell division, immune responses, morphogenesis of organs, and anatomical structures and cellular differentiation.

Among the stromal lineage cells, there is a strong tendency to intralineage communications, in which the interactions between the wnt5a ligands with receptors of the Ror1/2 and Fzd5a play a very important role. Activating the Fzd1 and Ror1/2 by Wnt5, triggers pathways associated with cell proliferation, differentiation, polarity, and apoptosis and have a positive effect on inflammatory diseases [21]–[23].In this network, the FRC.MLN stromal cell of mesenteric lymph nodes, has 32 loops with the most autocrine communication pathways, of which 32 communication pathways are only due to two interacting receptor-ligands, one of which is the interaction between the angiotensin-converting peptide hormone and its specific receptor. Macrophages exist in various tissues such as brain, lung, liver, spleen, bone marrow and lymph nodes. This cells are involved in immune and anti-tumoral responses to pathogens, maintenance of tissue homeostasis, and tissue repair [24], [25].

According to the results, the stromal cells have the most communication with other cells especially macrophages *(Fig. 3)*. In addition, bone marrow stromal cells play an important role in differentiating these cells by creating a suitable environment for growth and differentiation of hematopoietic cells. One of the most important immune cells that forms such an environment is macrophages [26]. Therefore, one of the reasons that the stromal cells in the network have the most communication with macrophages can be the role of these two cells in the process of hematopoietic cell differentiation.

The specificity of the expression of receptor and ligand genes and, consequently, the interactions they have with each other, can be attributed to the fact that the transmission of information between cells in a porous tissue is not a random process without planning. Therefore, the coordinated behavior of cells to maintain the stable state of multicellular organisms, which is largely dependent on intercellular communication, is that the transmission of messages between cells is selective. This is possible only if the expression of the components involved in the transmission of the message is specific to the type of sender and recipient cells [12], [27]

One of the most challenging steps in this study was the choice of the type of data needed to determine the receptor and the ligand used in each of our cells, such that, selection of each causes loss of some information needed for thorough assessment of this process. For example, the use of transcriptome data or proteomics information needs to ignore some receptor and ligand interactions, where the ligand molecule is non-protein such as steroid hormones. Similarly, the use of metabolomics data requires the exclusion of molecules from proteins. On the other hand, data on the interactions between metabolites and proteins are very low in the database.

One of the ways to overcome this limitation can be the simultaneous use of omics data such as transcriptomics, proteomics, and metabolomics which have been derived from a cell. By combining several omics data, complex biological processes such as intercellular communication can be more effectively analyzed. Needless to say, due to the lack of data obtained from the simultaneous analysis of several omics, which are also referred to as “multiomics analysis”, the use of this method to restructure the network is subject to limitations [28].

## Methods

The intercellular interaction network was reconstructed in two stages of data extraction and, integration and filtering of data. *Figure 7* shows the general stages of network reconstruction.

**Figure 7:**
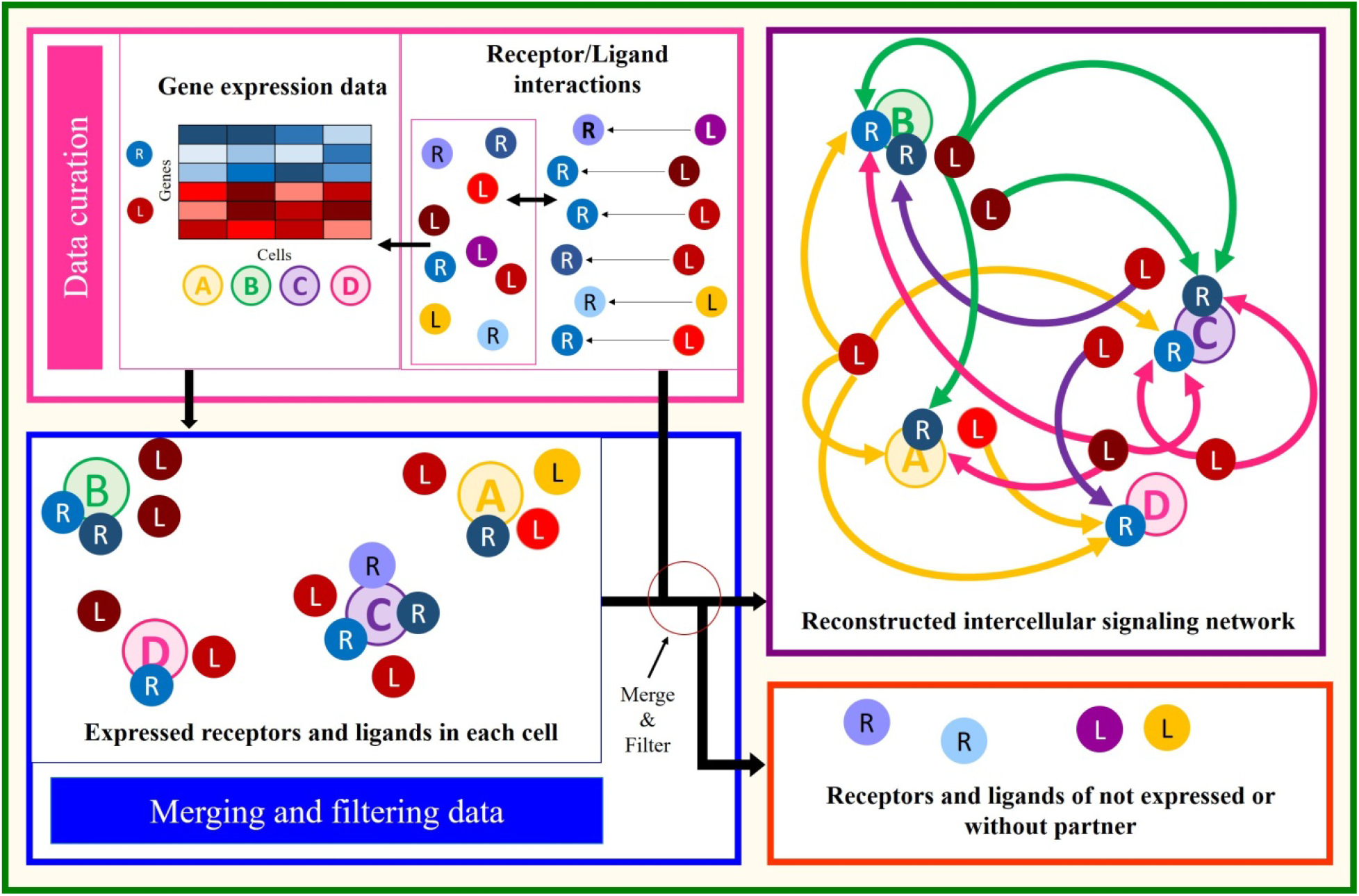
Schematic workflow for intercellular interaction network reconstruction.

### • Receptor–ligand interaction dataset

Receptor and ligand genes were extracted from the molecular databases such as the KEGG, GPCRdb, Uniprot, and IUPHAR [29]–[32]. In order to detect receptor-ligand interactions in the proteins interactions databases, the iRefWeb interface [33] was used and the receptor-ligand interactions (receptor-ligand pairs) among protein-protein interactions of the mouse organism were extracted. The data obtained from this source was added to the receptor and ligand interactions of previous related articles [34], [35] and IUPHAR and KEGG database. After removing the repetitive interactions, the benchmark dataset was created, which included 958 interactions between 340 receptors and 374 ligands.

### • Receptor-ligand gene expression dataset

The microarray expression data deposit in the Gene Expression Omnibus [36] (http://www.ncbi.nlm.nih.gov/geo/) under accession number GSE15907, which is related to the first phase of the immunological genomeproject (http://www.immgen.org/Protocols/ImmGen) [37]. The samples studied in this project (ImmGen) are from 24 different tissues of adult male B6 mouse. The Affymetrix 1.0 ST MuGene arrays platform was used to profile the expression of genes, based on criteria such as sensitivity, noise, different expression and reliability or diagnostic validity for each platform, using a collection of the common RNAs of two CD4^+^ and CD19^+^ cells, selected from among the several platforms used in the microarray analysis. This dataset consists of 653 samples related to 212 cell types, which include all major hematopoietic lineages [37].

These lineages are divided into 8 main lineages based on the hematopoietic lineage tree created in this project. The eight main hematopoietic lineages include stem and progenitor cell (S&P), granulocytes, monocytes, macrophages (MF), dendritic cells (DC), natural killer cells (NK), B cell and T cells. The T cells are divided into four subgroups of gamma and delta T cells (γδT), alpha and beta T cells (αβT), regulatory T cells (T_regs_) and natural killer T cells (NKT).In addition to the hematopoietic lineages, the stromal cells which are categorized as non-hematopoietic cells lineage, exist in this project [38].

from the 653 samples examined in the first phase of ImmGene immune Project, 25 samples of CD4^+^ and CD19^+^ cells were used as control and test samples to select the type of platform used for fine-array analysis of these samples, and the rest of the samples included 29 embryonic and 599 adult samples [37]. For analyzing this dataset, R-software and limma package were used and the method of normalizing the mean of several precise arrays (RMA) was used to normalize the data [39], [40].

After normalizing the data, Ensemble Biomart tool was used to map the probeset identifier to gene names [40], [41]. After the removal of probesets lacking gene names and probesets with more than one gene name [42], a collection of 25751 probesets, each mapped to a single gene name, was created. For 3726 gene names that had more than one probeset on the arrays, the probeset identifier was assigned to the gene name which had the highest amount of expression between the probesets [38].

Subsequently, using the receptor-ligand interactions dataset, the probesets belonging to the genes encoding receptor and ligand proteins were extracted from this collection *(Fig. 8 a)*. After identifying the genes of the receptor and ligand in each cell sample from the threshold defined in equation 1, which represented the expressed genes of receptors and ligands, was used for each cell. Because there was more than one sample from each cell type, before the threshold was applied, the average expression of the same cell samples was obtained and finally the threshold was applied on 199 samples which were representatives of each cell type. In order to reconstruct the network, those genes whose according to the threshold defined in equation 1, expressions were above their mean expression (X_i_) plus three times the standard deviation of their expression (σi) in all samples, was considered as the expressed genes in each cell [13]. If the expression level of each gene is greater than the defined threshold, then the number (1) and otherwise (0) are assigned to them. *Figure 8b* shows how to obtain the expressed genes of the receptor and the ligand in 199 cell samples.

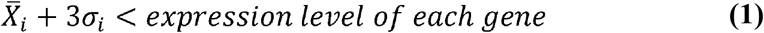

**Figure 8:**
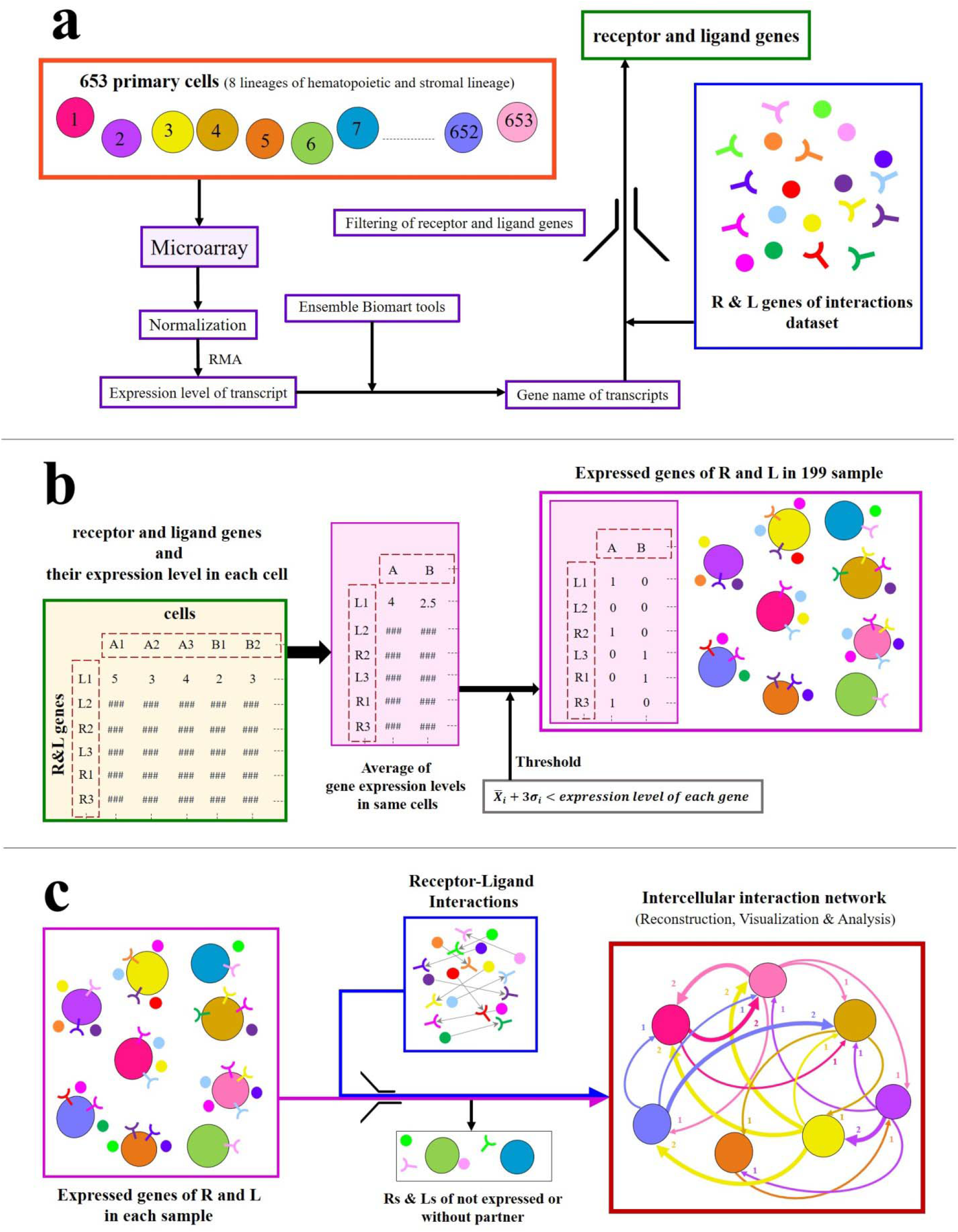
Creating a dataset of expressed receptor and ligand genes. **a)** Identification of receptor and ligand genes and their expression level in each sample. **b)** Determining the receptor and ligand genes expressed in each sample by applying the threshold. **c)** Reconstructing, visualizing and analyzing the intercellular interconnection network by integrating the receptor-ligand interaction dataset with the dataset of receptors and ligands expressed in each cell sample.

### • Reconstruction of intercellular communications

According to the dataset of the receptor-ligand interactions and the list of receptor-ligand genes expressed in all the samples, only those receptor-ligand pairs or interactions were selected in which the expression of both the ligand and receptor was above the defined threshold (equation 1). By identifying these interactions, communication between the cells is also indicated. For example, if the expression of a pair of receptor-ligands in two cells is higher than the specified threshold, obviously those two cells, one of which expresses the ligand gene and the other receptor gene, have intercellular communication with one another [13], [43]. Next, the network reconstructed interactions were visualized and analyzed by open source software of Cytoscape *(Fig. 8 c)* [44].

GEO: Gene Expression Omnibus ()
ImmGen: Immunological genome project
S&P: stem and progenitor cell
MF: macrophages
DC: dendritic cells
NK: natural killer cells
γδT: gamma and delta T cells
αßT: alpha and beta T cells
T_regs:_: regulatory T cells
NKT: natural killer T cells
RMA: several precise arrays
LNs: lymph nodes

